# A new interesting and unidentified Schmidtea species in Romania (Platyhelminthes, Tricladida, Dugesiidae)

**DOI:** 10.1101/2024.02.26.582039

**Authors:** Anda Felicia Babalean

## Abstract

This paper presents the characters (morphology and aspects of reproductive biology) of a population belonging to the genus *Schmidtea*, in Romania. This population shows similarities with the other Schmidtea species: with *S. polychroa*, the presence of cephalic sensory fossae; with *S. mediterranea*, the asexual reproduction by fission; with *S. nova*, the presence of a diaphragm of the ejaculatory duct to the secondary seminal vesicle. Set of characters make the population distinct and thus, candidate for a new species: the disposition / arrangement of the genital atria; a wide and straight / narrow and convoluted ejaculatory duct within the penis papilla; in some specimens the penis papilla has two lobes, each with own duct.

## 1. Introduction and historical background

Schmidtea Ball, 1974 is a genus of a freshwater flatworm with a geographical range restricted in the West Palearctic region (Leria et al. 2018). It was introduced in the systematic of the Family Dugesiidae at the genus rank based on two morphological characters of the copulatory apparatus: the penis bulb with a double seminal vesicle, and the mixed musculature of the bursal canal (de Vries & Sluys 1991). The genus Schmidtea is considered monophyletic both on morphological and molecular data (Sluys 2001; Riutort et al. 1992).

After a tangled taxonomic history, it was established that the genus Schmidtea includes four species: *S. polychroa* (Schmidt, 1861), *S. lugubris* (Schmidt, 1861), *S. mediterranea* (Benazzi, Baguñà, Ballester, Puccinelli, Del Papa, 1975) and *S. nova* (Bennazi, 1982) (Leria et al. 2018). The latter species was redescribed by Leria et al. (2018). It is appreciated that *S. lugubris* and *S. nova* are poorly studied (Leria et al. 2018).

In Romania there were recorded *S. lugubris* (Năstăsescu 1973, Leria et al. 2018) and *S. nova* (Leria et al. 2018). Another potential new Schmidtea species, showing numerous similarities with *S. mediterranea* was recorded by Babalean (2022).

A.F. Babalean dedicates this study to the outstanding scientists Kendra Greenlee, Axel L. Schönhofer, DW and DW.

## 2. Material and method

### The study is based on the following samples

**05 Nobember 2020**: 26 specimens fixed in Beauchamp solution for 2 hours thereafter, removed and stored in 70-75 ethanol. Two of the 26 specimens were prepared for usual histological serial sagittal sections (specimen Fl1 – 42 slides) and frontal (horizontal) sections (specimen Fl2 – 12 slides). They were embedded in paraffin wax, sectioned at 5 μm, stained in Haematoxilin-Eosin).

**05 July 2021**: 27 pieces (whole worms and fragments resulted from fission) fixed in Beauchamp solution, 2 specimens in 96 ethanol.

**07 March 2022**: 10 specimens fixed in Beauchamp solution, 12 specimens fixed in 96 ethanol (removed and stored in 70 ethanol), 1 specimen in absolute ethanol (?).

## 3. Results

### 3.1 Systematic

Order Tricladida Lang, 1881

Suborder Continenticola Carranza, Littlewood, Clough, Ruiz-Trillo, Baguñà, Riutort, 1998 Genus Schmidtea Ball, 1974

*Schmidtea species*

### 3.2 Morphology

#### *The whole specimens* – Figs. 1, 2

Living specimens measure up to 18-19 mm long – Fig. 1. Preserved specimens measure between 2,5 – 8,5 mm long (sample 05 July 2021) and between 5 – 10 mm (sample 07 March 2022) – Fig. 2. Colour blackish / black dorsally, dark brown to black ventrally. Head obtusely pointed or rounded, auricular region barely marked, neck region indistinctive. Two eyes set in pigment-free patches, placed very closed to the anterior margin of the head. Numerous specimens and particularly heads resulted from fission show only the pigment-free patches, yet no pigmented eye. One specimen with 4 eyes (supernumerary eyes). The ventral margin of the head presents whitish and luminous spots which are interpreted as sensory fossae – Fig. 2.

**Fig. 1.**
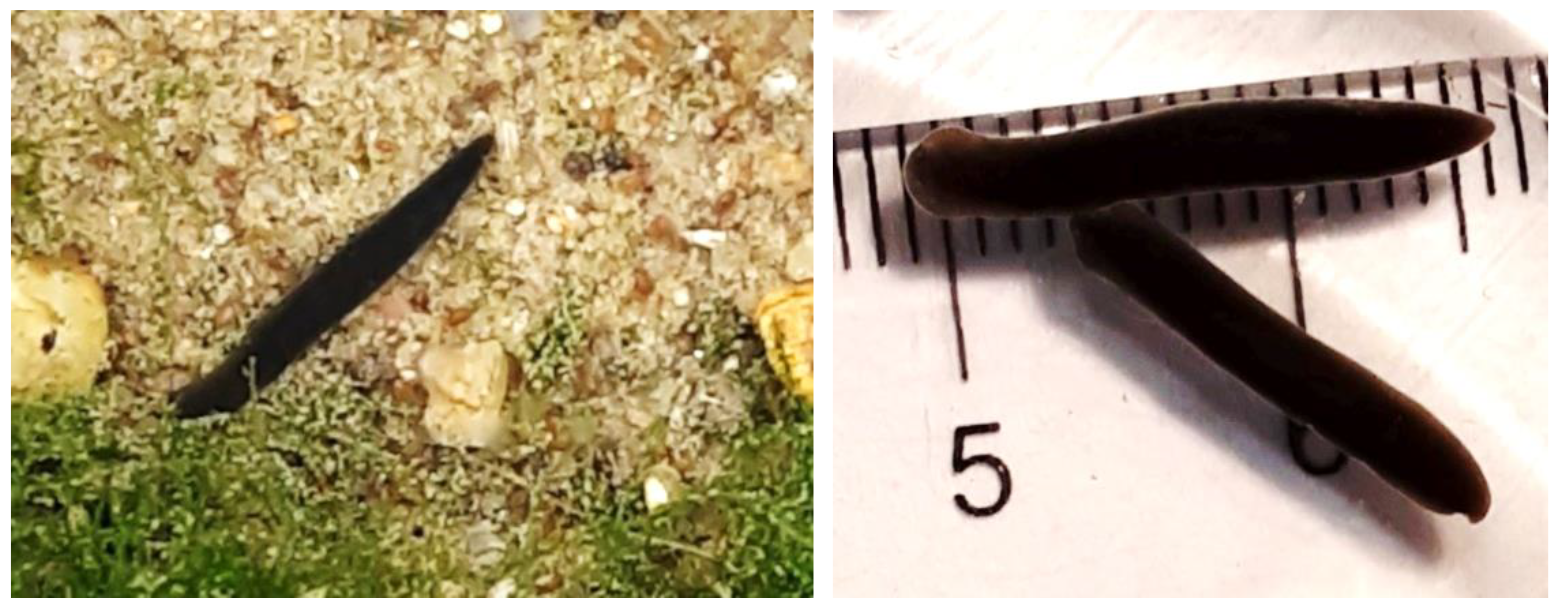
Living specimens in nature (left) and in laboratory (right)

**Fig. 2.**
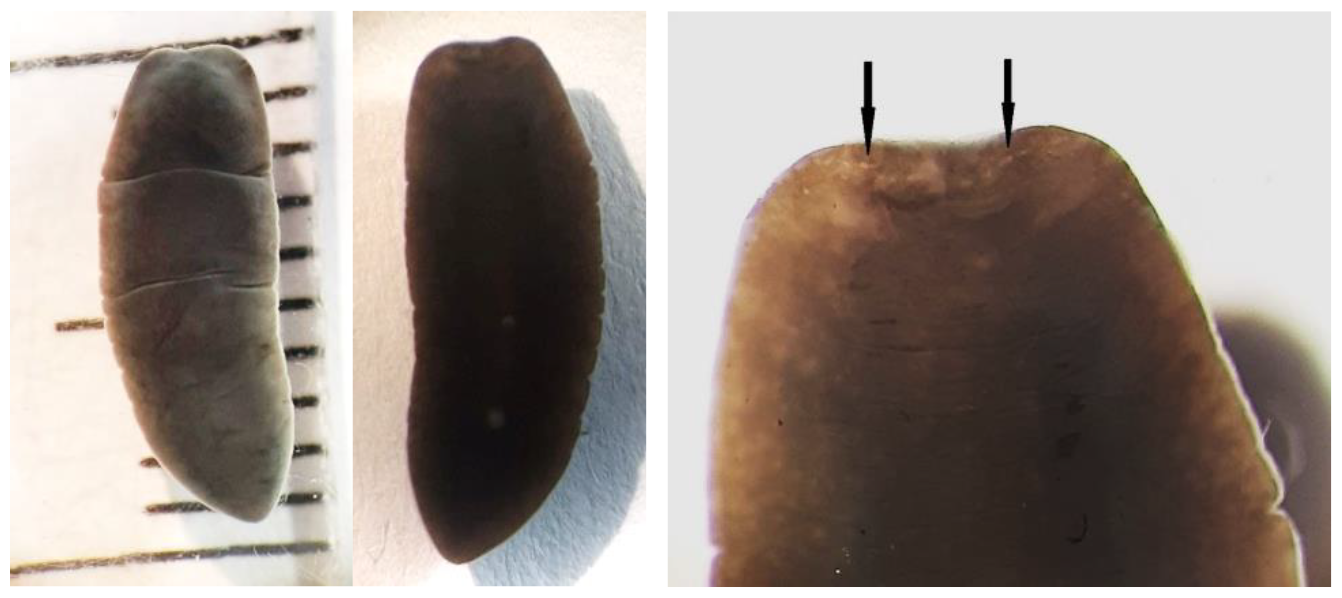
A fixed specimen on dorsal and ventral, showing sensory fossae (arrows)

Pharynx short and located in the second half of the body; the musculature of the planariid type, cf. Sluys (1989, Fig. 2)

Numerous specimens, especially those fixed in ethanol show the penis papilla extruded / protruded (it is assumed that the penis bulb cannot be protruded, being enclosed into the parenchymal tissue). The protruded papillae vary in length and shape, some are long and straight, some are short, some present an annular swelling just below the tip – Fig. 3.

**Fig. 3.**
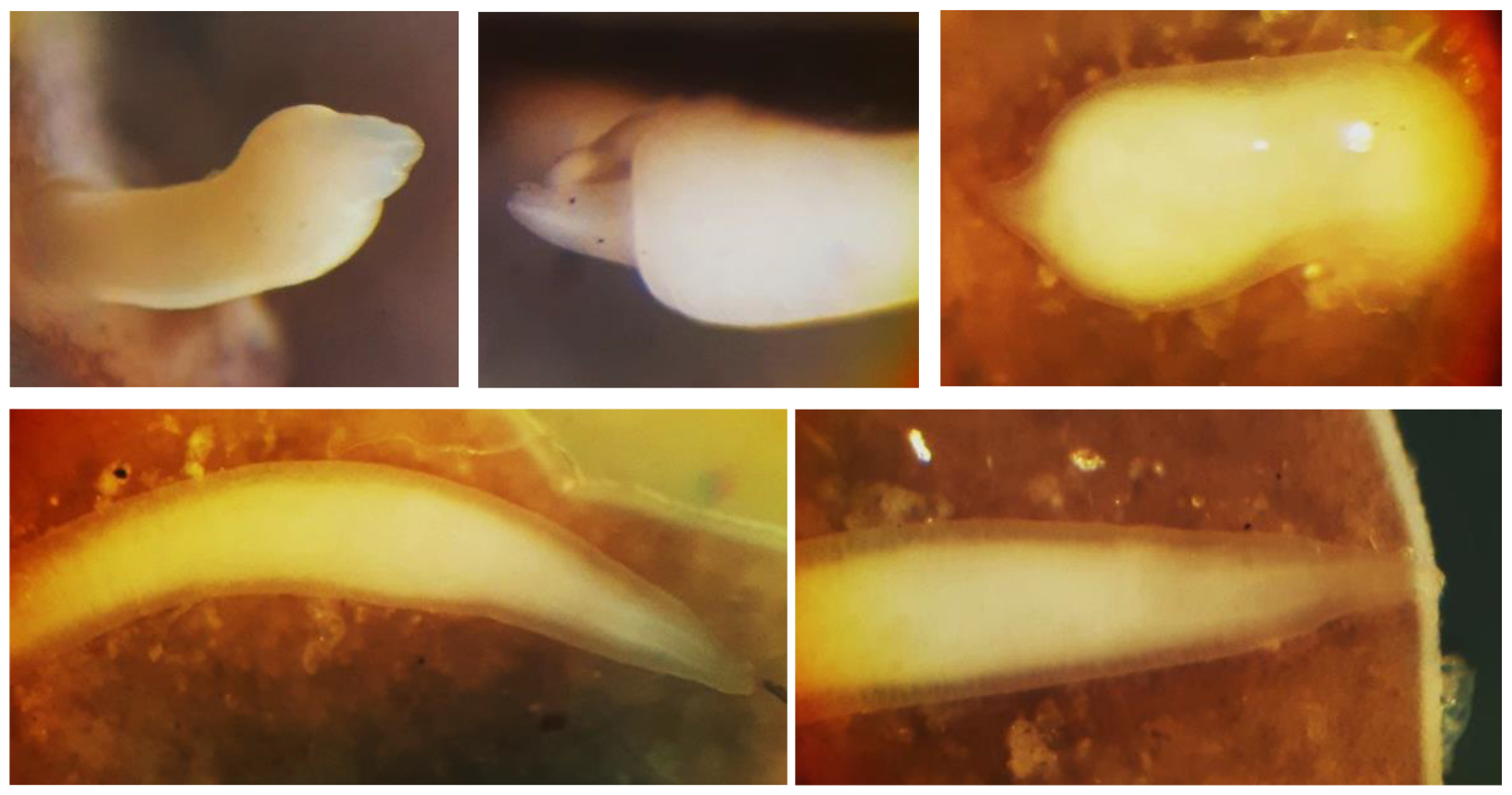
Aspect of the extruded penis papilla in some specimens: up – I1 (07 March 2022), I3 (07 March 2023), I5 (05 July 2021); down – I5 (05 July 2021), I1 (05 July 2021)

Follicular testes (present in both specimens Fl1 and Fl2) situated dorsally, on lateral sides of the body, throughout the body length. Ventral ovaries present in both specimens.

The gross anatomy of the copulatory apparatus – Figs. 4 – 16

**Fig. 4.**
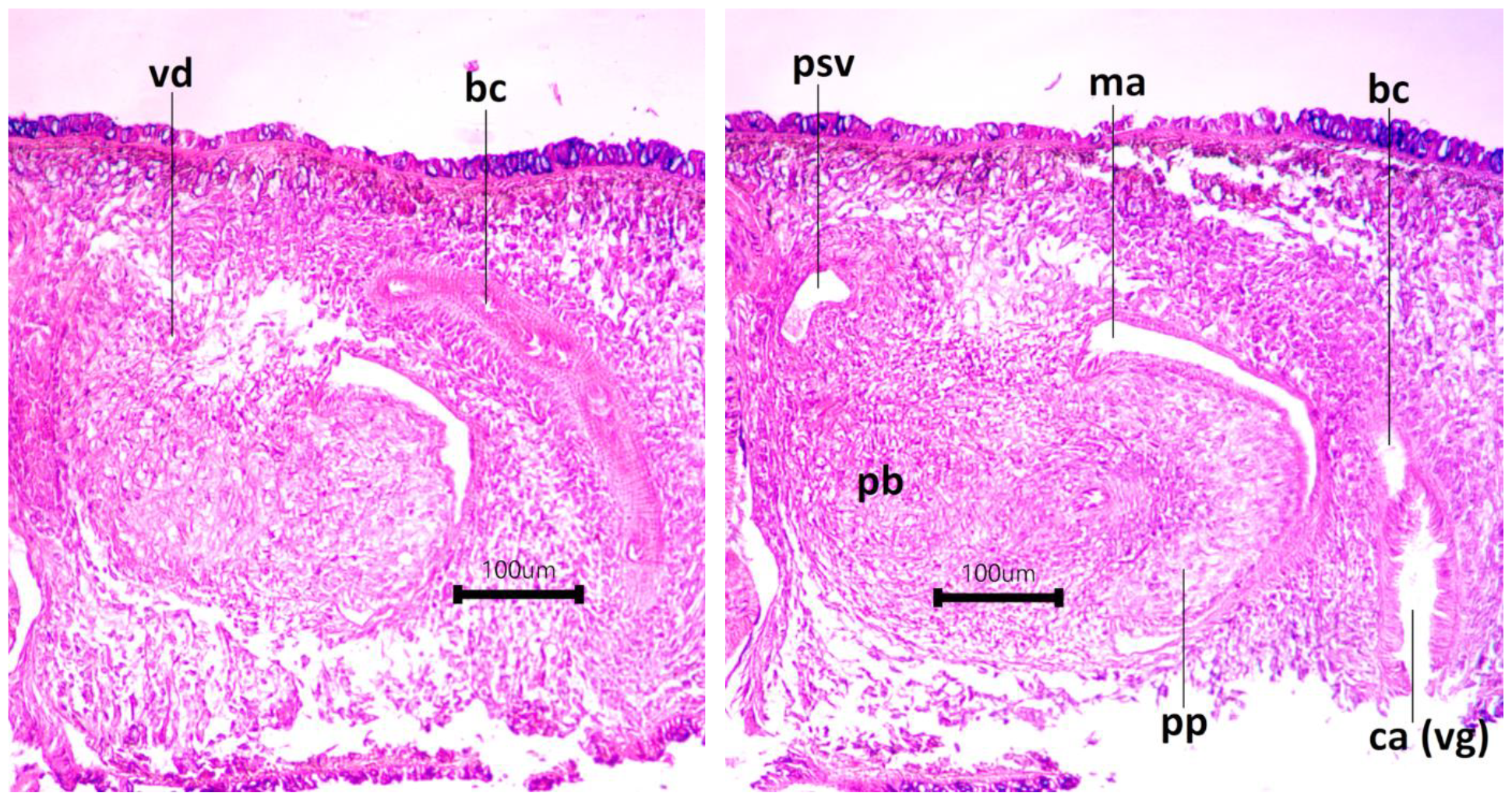
Microphotographs of the copulatory apparatus in sagittal sections, slides Fl1-23-5 (left) and Fl1-26-1 (right)

Examined on sagittal and frontal histological sections, the copulatory apparatus shows the following features:

- the vasa deferentia covered by a cuboidal epithelium, form very large spermiducal vesicles at the level of the pharynx, visible only in frontal sections (specimen Fl2) – Figs. 11, 12, 13.

- the penis bulb is a very well-developed muscular mass, consisting of two regions, each with own seminal vesicle. Thus, the penis bulb houses a double seminal vesicle: i) an anterior primary seminal vesicle (psv) surrounded by a wall of an internal thin layer of circular musculature and an external layer of irregular musculature, and ii) the posterior secondary seminal vesicle (ssv) surrounded by a thicker wall consisting of especially concentric layers of musculature – Figs. 4, 6, 8, 9, 15. The two regions of the penis bulb are interconnected by a muscular constriction with a narrow duct (lumen) which is part of the ejaculatory duct. It opens through a diaphragm into the secondary seminal vesicle (ssv) as revealed by the slide Fl1.27.4 – Figs. 4, 5, 6.

**Fig. 5.**
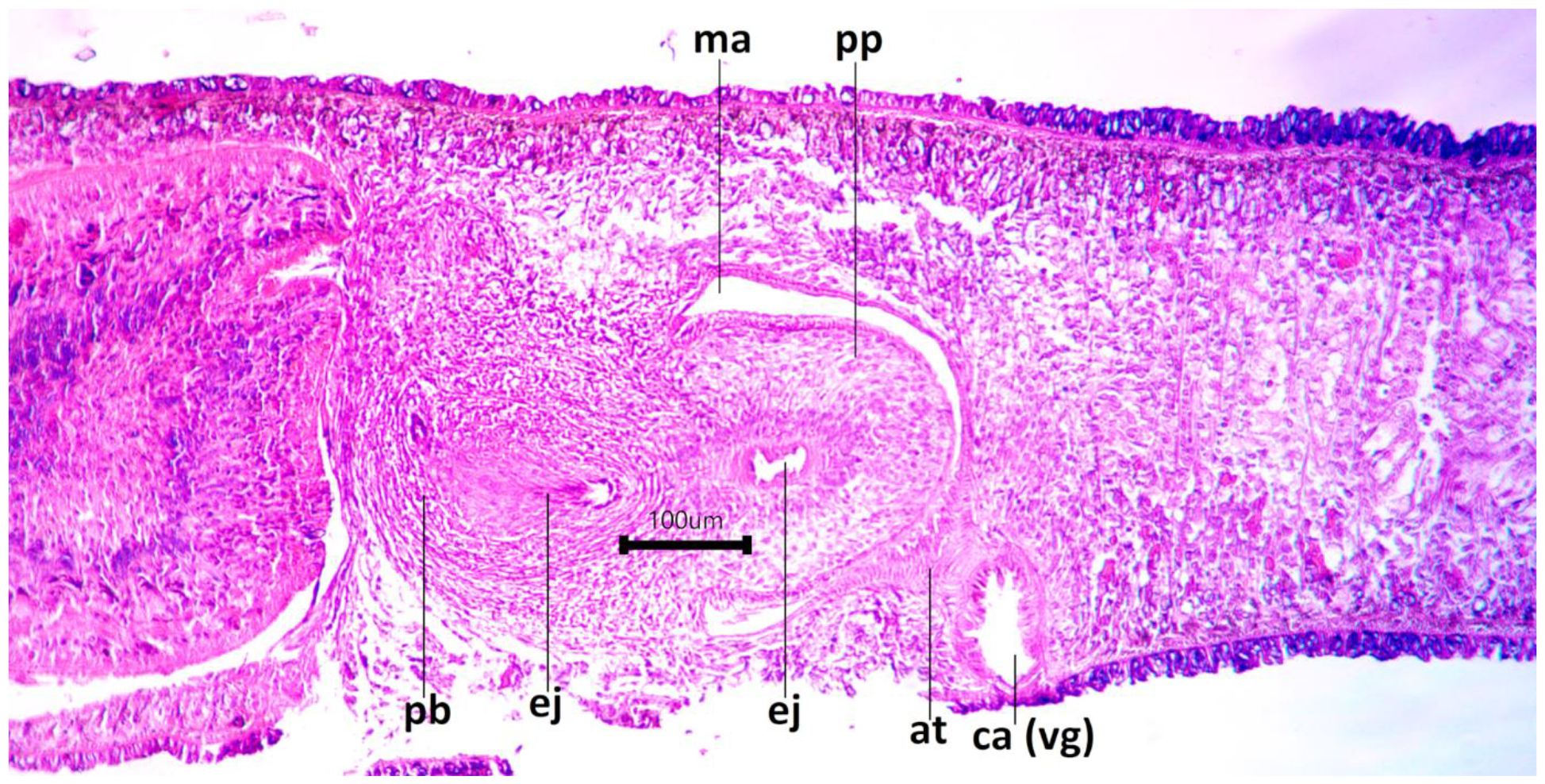
Microphotograph of the copulatory apparatus in sagittal sections, slide Fl1-27-2

**Fig. 6.**
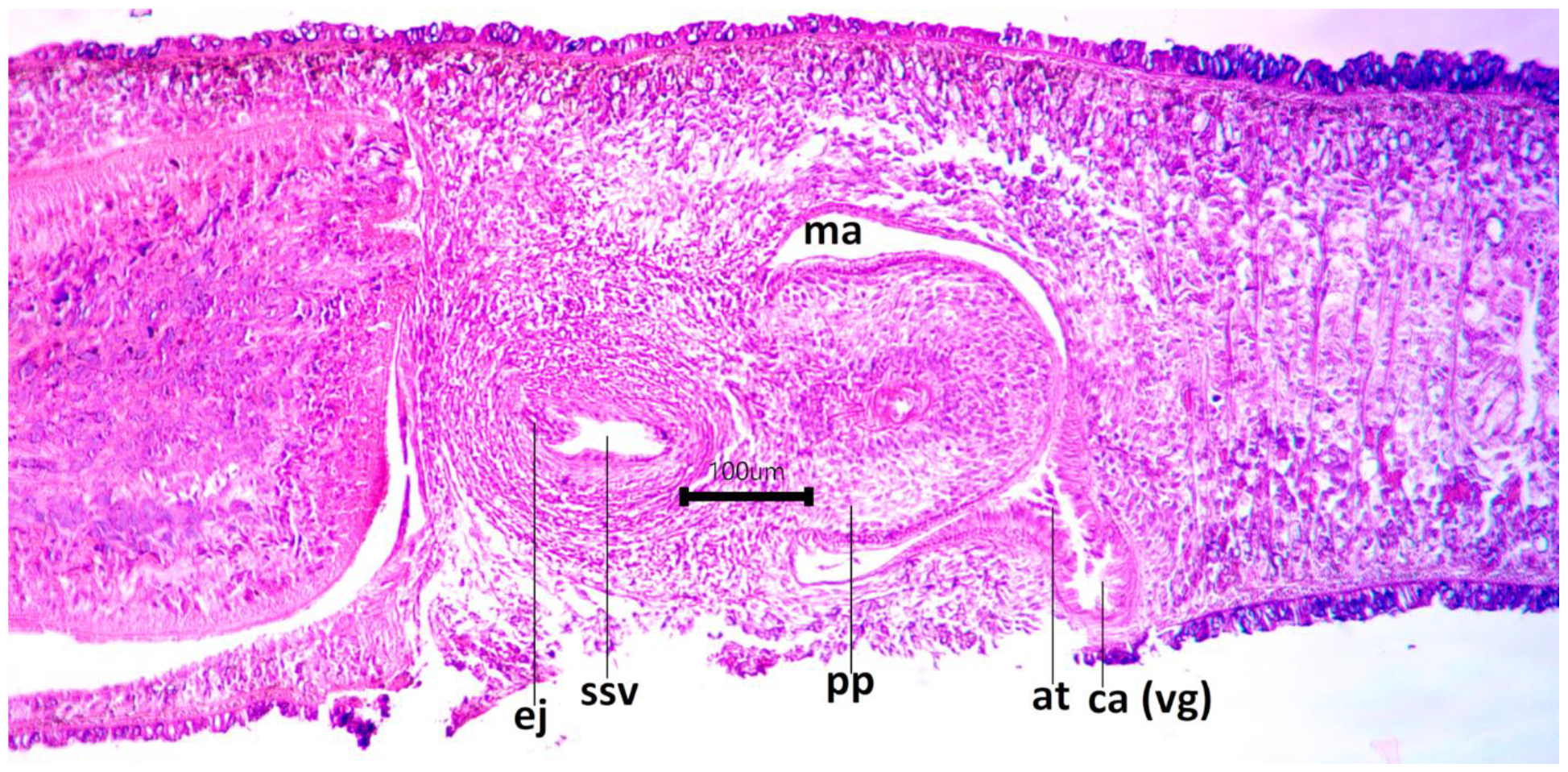
Microphotograph of the copulatory apparatus in sagittal sections, slide Fl1-27-4

- the vasa deferentia present a 90-degree curvature before they open separately into the primary seminal vesicle – Figs 9, 10.

- the penis papilla appears to be very short in both the sagittal and frontal sections. It contains internal cellular tissue. The penis papilla occupies the male genital atrium which is clearly separated by the common hermaphroditic genital atrium. The male genital atrium is extended with an atrial tube (at) which opens into a small ventral cavity which may be considered the common genital atrium (ca) or the vaginal area of the bursal canal (vg) – Figs. 5-8, 11-13, 15.

- the ejaculatory duct within the penis papilla is very wide in the sagittal sections – Figs. 5, 7, 8. In the frontal sections, the aspect of the ejaculatory duct is rather narrow and convoluted, included in a wide space filled with a pink mass – Figs. 14, 15.

**Fig. 7.**
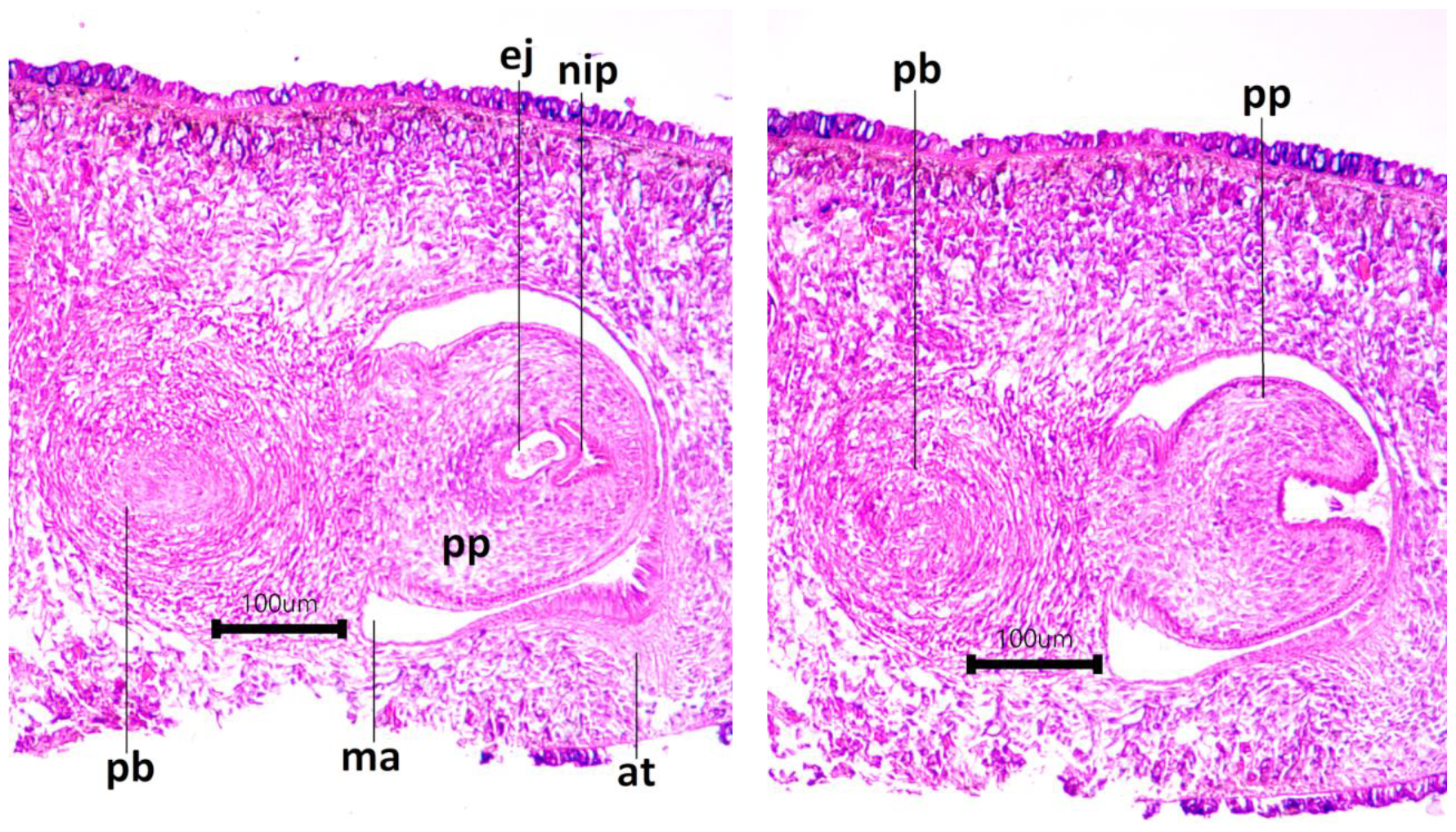
Microphotographs of the copulatory apparatus in sagittal sections, slides Fl1-28-2 (left) and Fl1-29-2 (right)

**Fig. 8.**
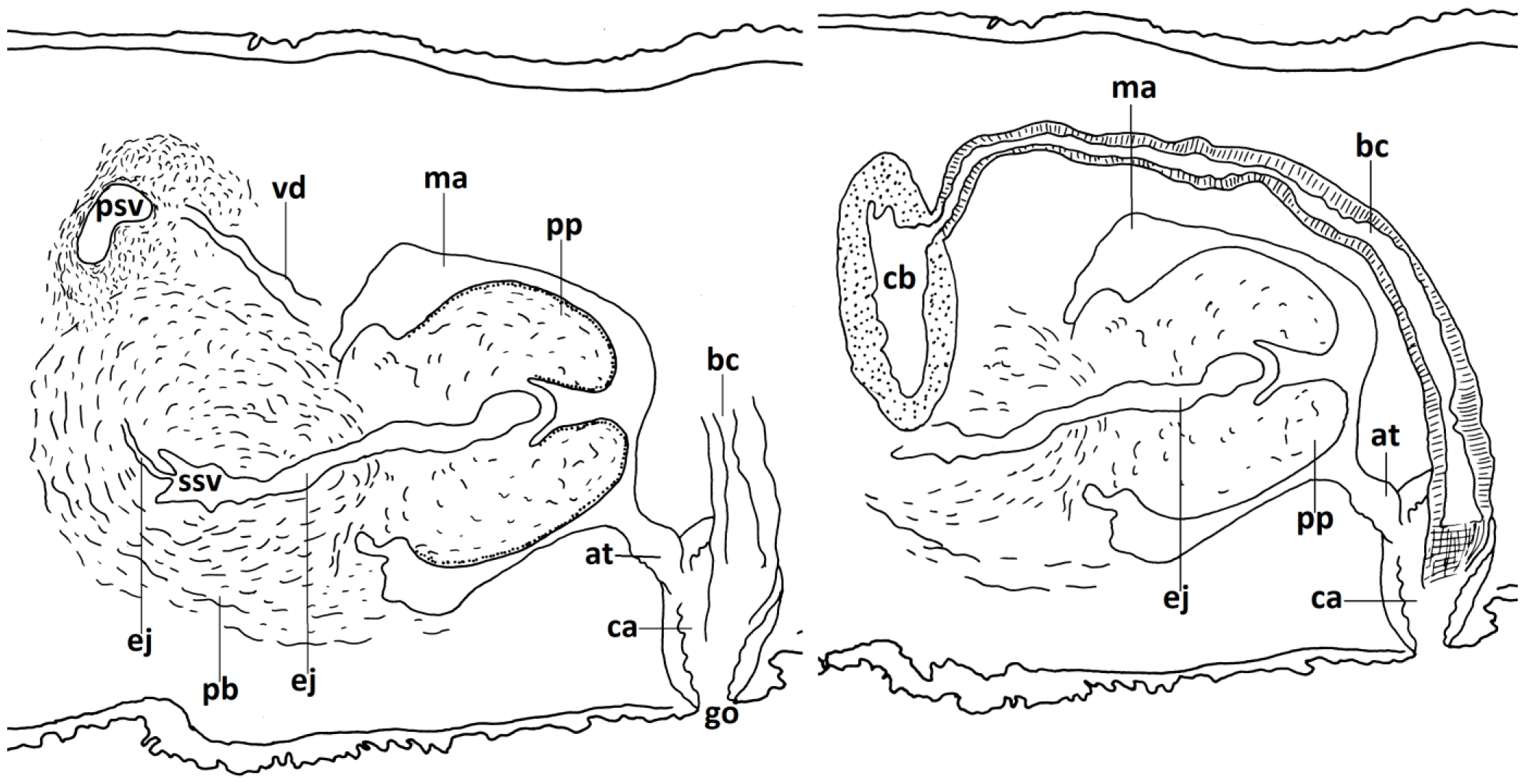
Diagrammatic reconstruction of the copulatory apparatus in sagittal section, specimen Fl1, male (left), female (right)

**Fig. 9.**
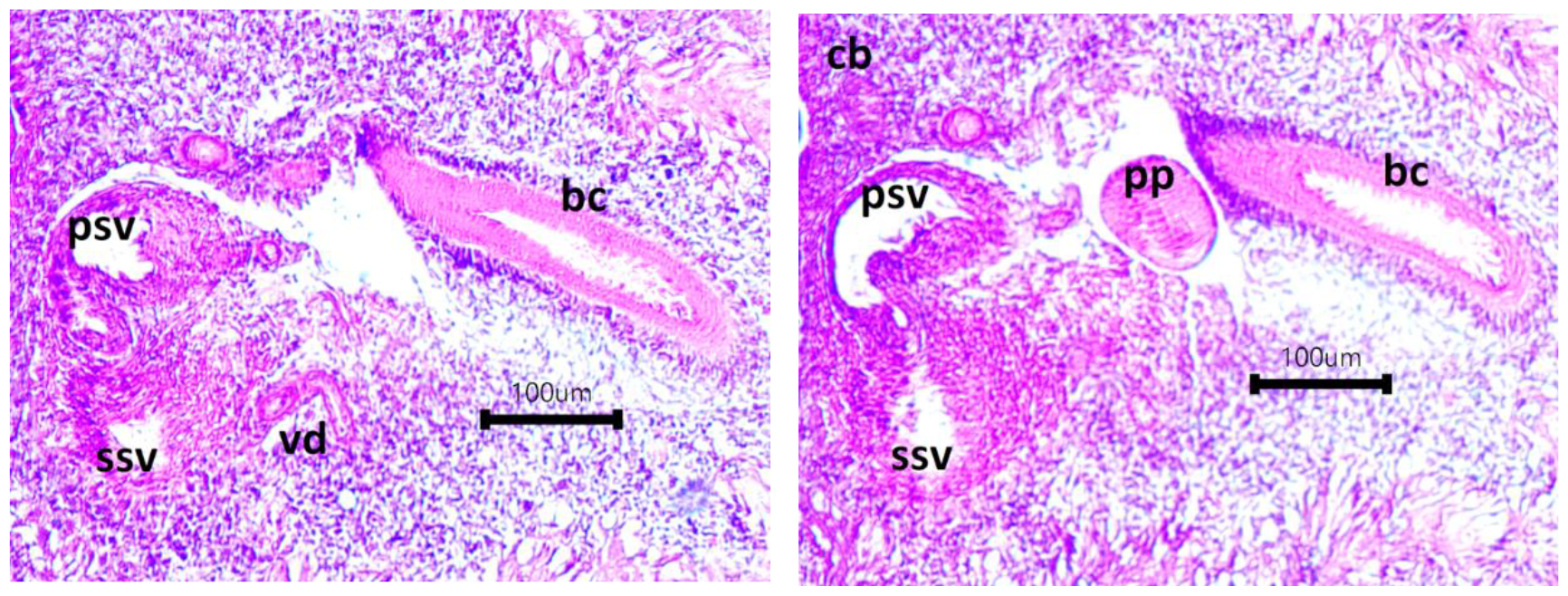
Microphotographs of the copulatory apparatus in frontal sections, slides Fl2-10-4 (left) and Fl2-10-1 (right)

**Fig. 10.**
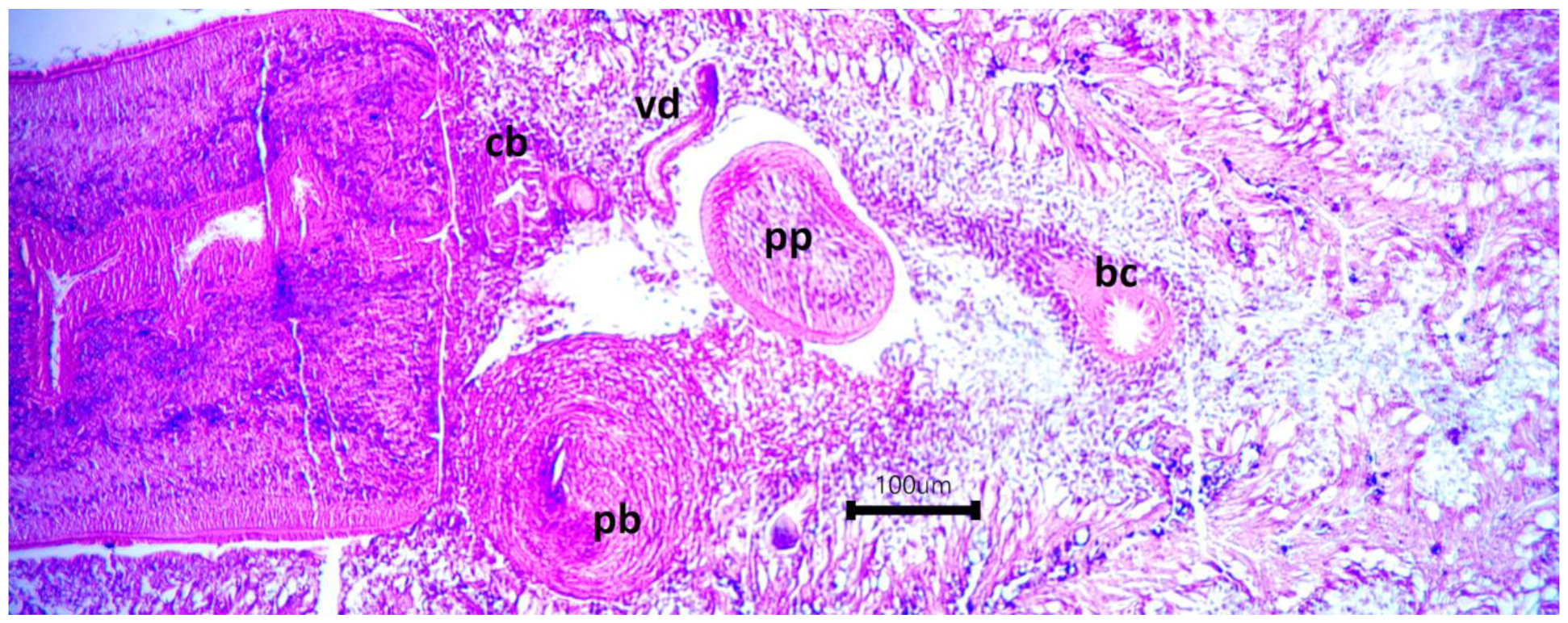
Microphotograph of the copulatory apparatus in frontal sections, slide Fl2-8-2

**Fig. 11.**
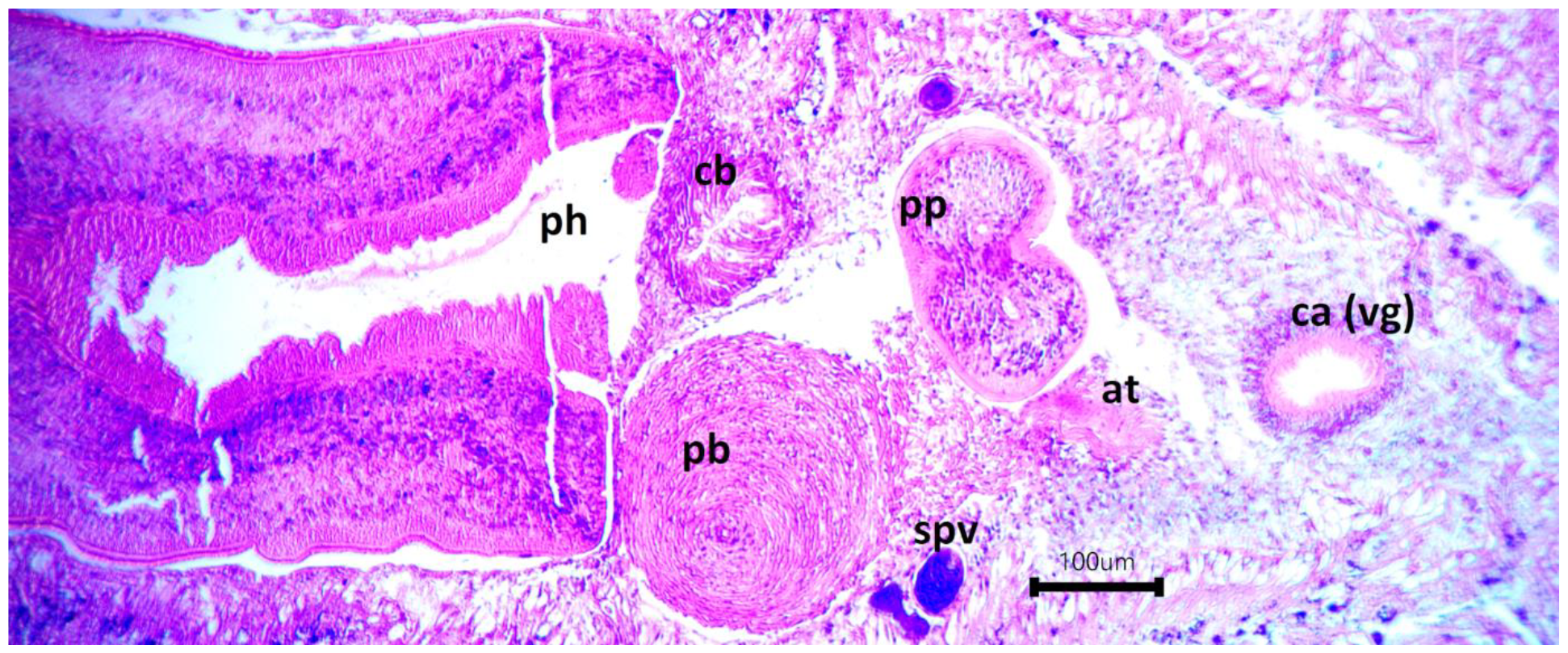
Microphotograph of the copulatory apparatus in frontal sections, slide Fl2-7-1, the penis papilla shows a strangulation that will give two lobes

**Fig. 12.**
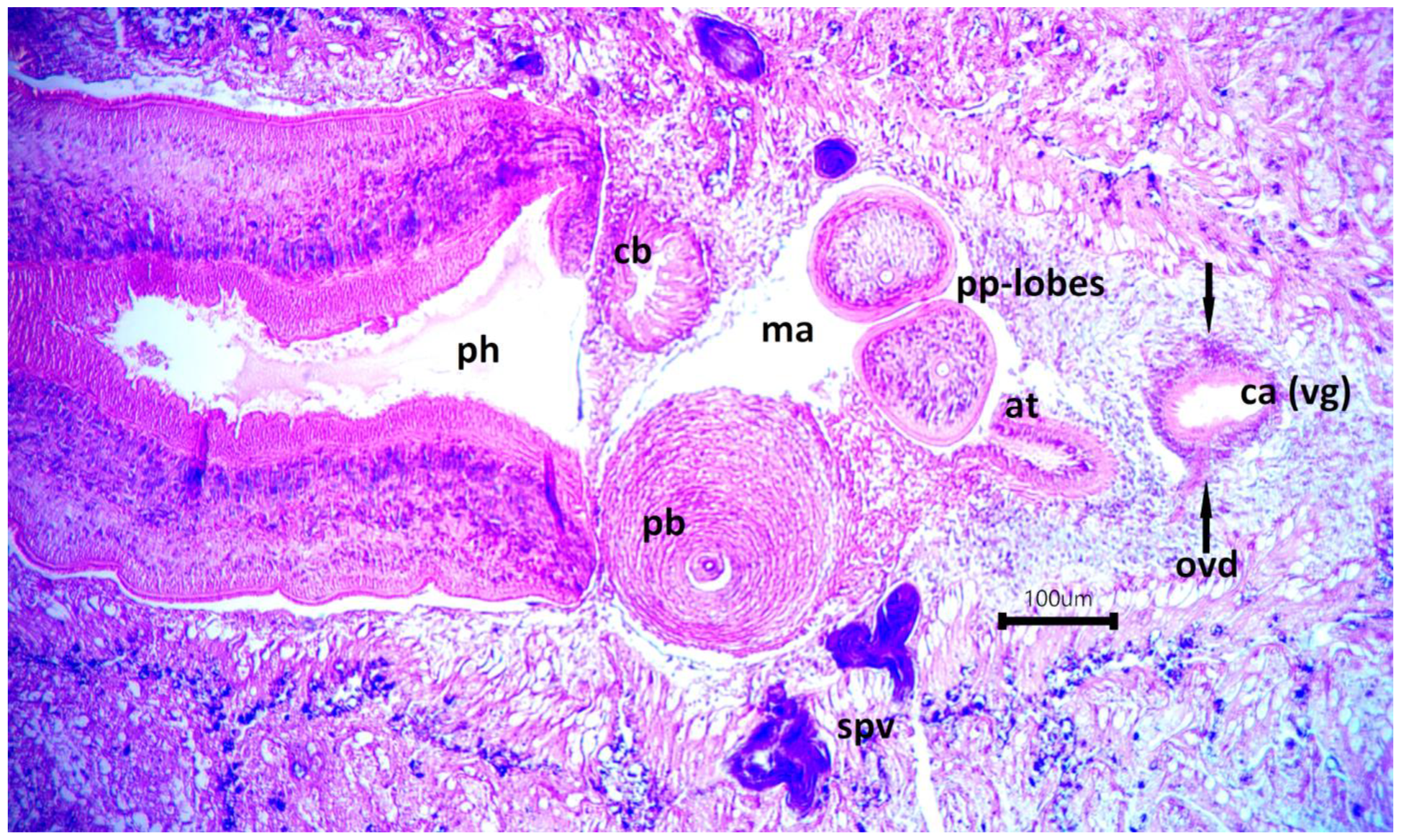
Microphotograph of the copulatory apparatus in frontal sections, slide Fl2-6-1

**Fig. 13.**
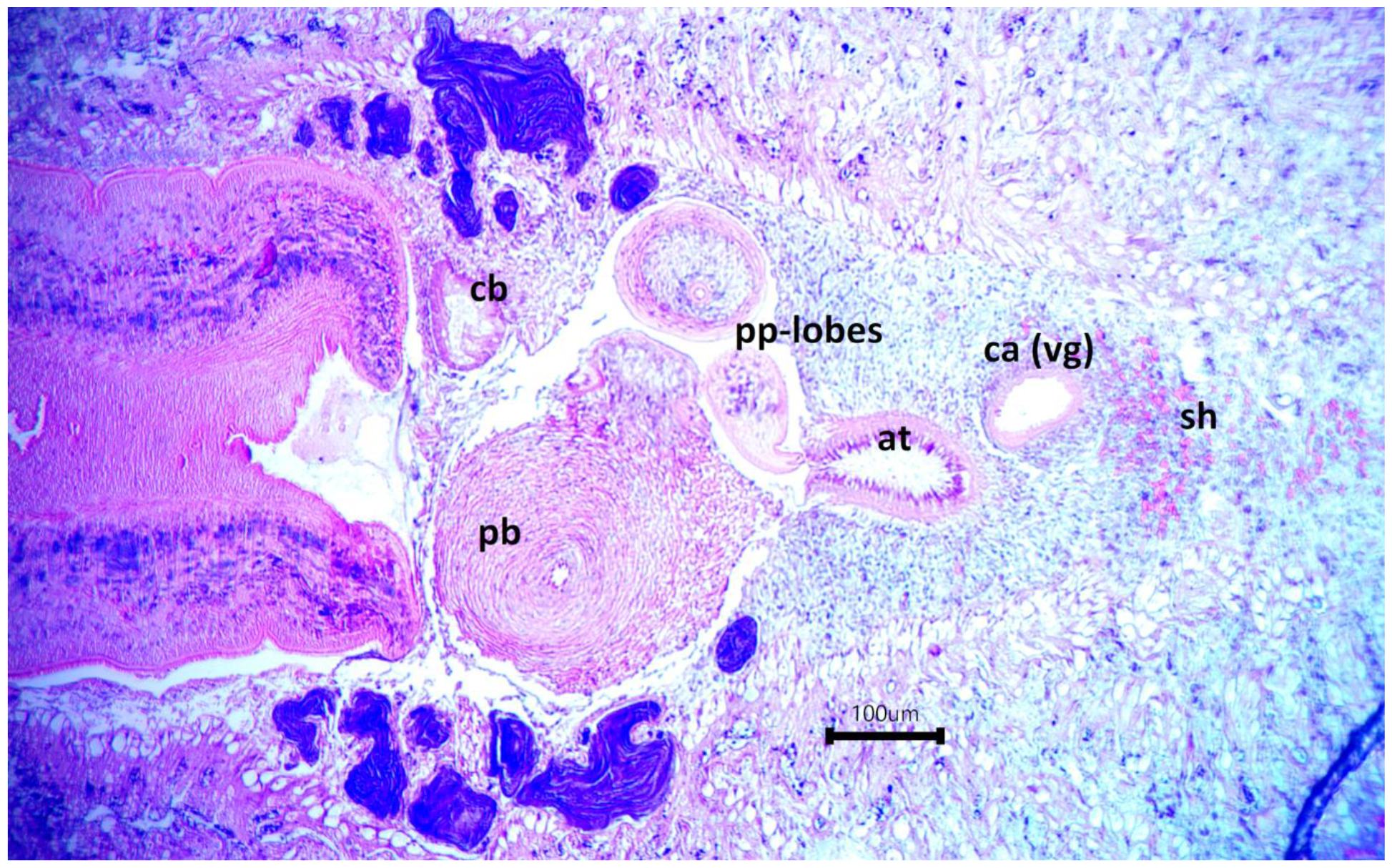
Microphotograph of the copulatory apparatus in frontal sections, slide Fl2-4-2

**Fig. 14.**
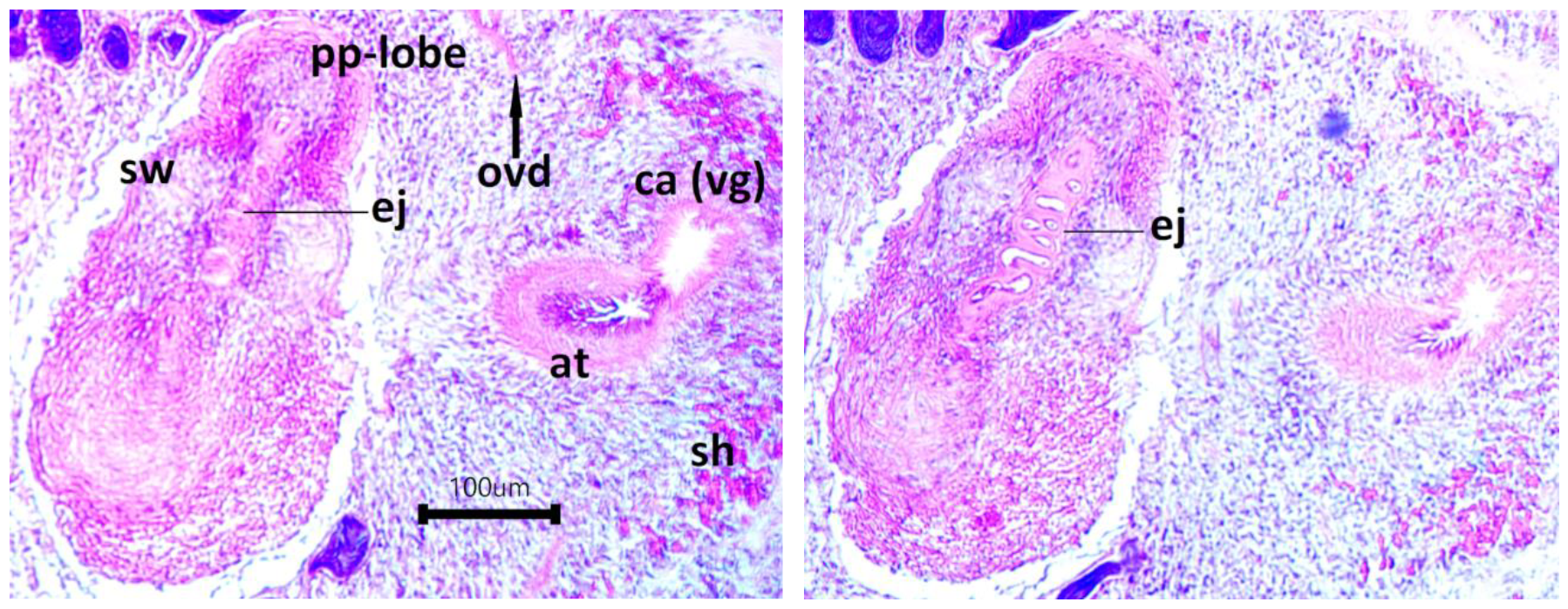
Microphotographs of the copulatory apparatus in frontal sections, slides Fl2-4-1 (left) and Fl2-3-4 (right)

**Fig. 15.**
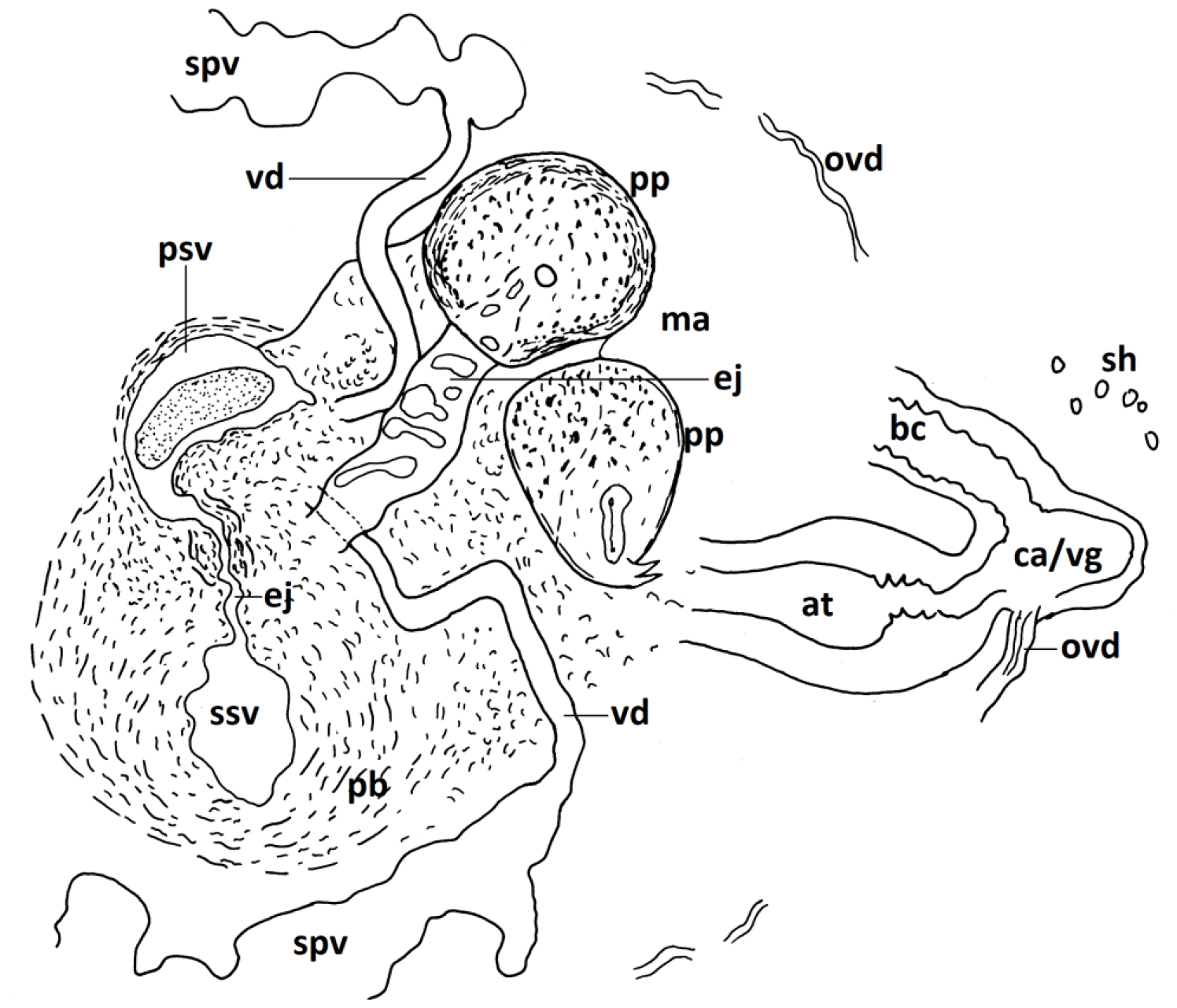
Diagrammatic reconstruction of the copulatory apparatus in frontal section, specimen Fl2, male part

- the length of the penis papillae (short or long, in histological slides, also the protruded ones) may be an individual character or may be the result of a relaxed or a contracted state, for instance a strong muscular contraction induced by work reagents or a relaxed, not-contracted physiological state.

- the penis papilla is bilobed in both sagittal and frontal sections. In inside, the lobes contain cells covered with layers of circular musculature – Figs. 12, 13. In the frontal sections (Fl2) the two lobes of the papilla show their own ducts – Figs. 12,13,15. Their pink coat indicates they come / originate from the wide space within the penis papilla before it becomes bilobed. Through semi-transparency, some protruding papillae leave the impression of a duct in each of the two lobes – Fig. 3.

- the copulatory bursa has a dorsal position, under the pharynx. The bursal canal has a very wide lumen surrounded by a wall of intermingled musculature cf. to DeVries & Sluys (1991, Fig. 4) – Figs. 4, 9. The bursal canal runs posteriorly over the penis, takes a descendent not-angled course coming posterior to the male atrium, thereafter, opens into the small ventral cavity considered either the common genital atrium or the vaginal area of the bursal canal. This small cavity opens to the outside through the gonopore.

- the two oviducts are visible only in the frontal sections and they appear to open separately into the common genital atrium or into the vaginal area of the bursal canal – Figs. 12, 14, 15, 16.

**Fig. 16.**
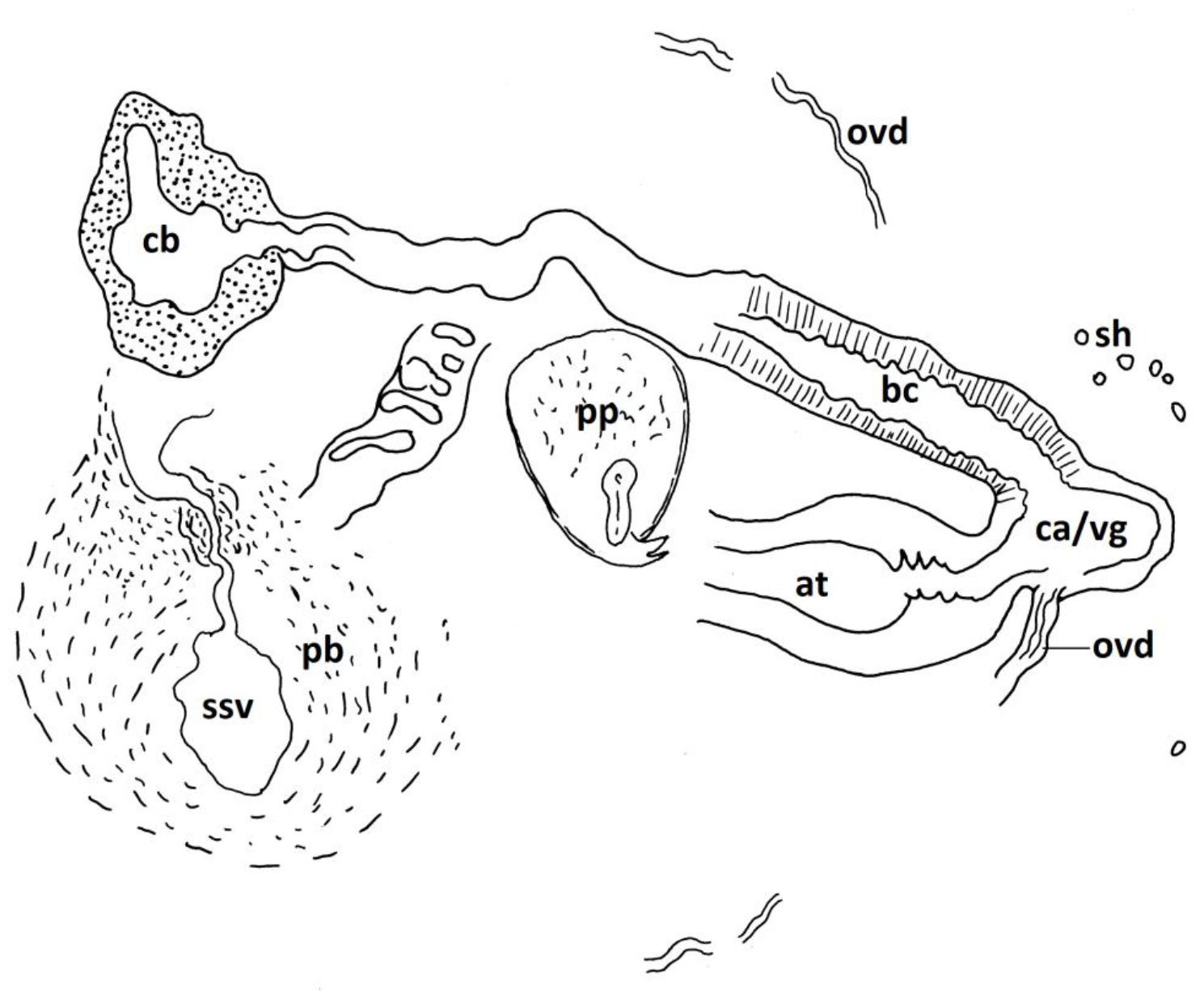
Diagrammatic reconstruction of the copulatory apparatus in frontal section, specimen Fl2, female part

#### *The fragments resulted from fission* – Fig. 17

The sample contains 6 pieces resulted from fission:

**Fig. 17.**
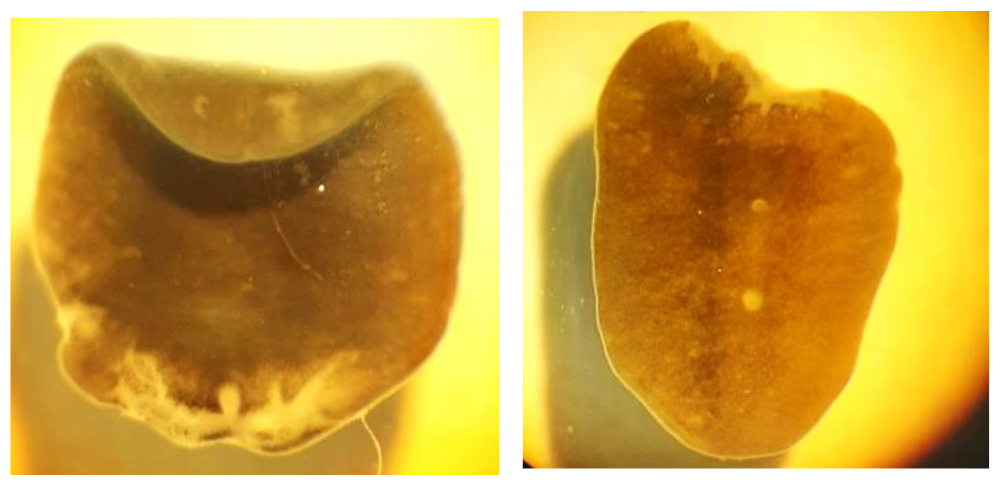
Fragments of worms resulted from fission

- only the head with eyes and no orifices (mouth and gonopore) – 3 pieces

- the body with both orifices (mouth and gonopore) – 2 pieces

- body with head and only one orifice (mouth) – 1 piece

### 3.3 Reproductive biology

Sexually mature worms were collected in November, March, July. Fission occurs in July.

## 4. Discussions

### Comparative morphology

The following characters are comparatively discussed:

#### 1) The penis bulb and internal cavities

The penis bulb with a well-developed muscular mass and with a double seminal vesicle (a primary seminal vesicle and a secondary seminal vesicle) is characteristic for the genus Schmidtea. The distinction between species concerns other aspects of the penis bulb: it is elongated in *S. lugubris* (Ball & Reynoldson 1981); “reflected dorsad” (with the primary seminal vesicle oriented dorsal to the secondary seminal vesicle) in *S. poychroa* (Ball & Reynoldson 1981), in *S. mediterranea* (Benazzi et al. 1975, Fig. 1; Harrath et al. 2004, Fig. 1), *S. nova* (Leria et al. 2018, Fig. 10), the specimens of this paper (Figs. 4, 8 sagittal Fl1). The two seminal vesicles (psv and ssv) are interconnected through a narrow ejaculatory duct which runs without any diaphragm in *S. lugubris, S. polychroa* (Ball & Reynoldson 1981) and in *S. mediterranea* (Benazzi et al. 1975, Harrath et al. 2004). The ejaculatory duct within the penis bulb shows a diaphragm to ssv in *S. nova* (Leria et al. 2018, Fig. 10) and in specimen Fl1 of this paper (Figs. 4, 8). A clear muscular constriction between the two parts of the penis bulb is visible in frontal sections in Fl2 specimen of this paper – Figs. 9, 15, 16.

#### 2) The penis papilla and its ejaculatory duct

The penis papilla is a character much more variable between species. It has a conical shape in *S. lugubris, S. polychroa*, longer in *S. lugubris*, shorter and with a dorsal hump in *S. polychroa*, with the ejaculatory duct running centrally in *S. lugubris*, and ventrally in *S. polychroa* (Ball & Reynoldson 1981). In *S. mediterranea*, the penis papilla is a long cone with the ejaculatory duct running centrally (Benazzi et al. 1975). The penis papilla of *S. nova* is distinct by a knee-shaped bend and the wide ejaculatory duct in its terminal part, thus giving the aspect of two terminal lobes (Leria et al. 2018, Fig. 10-11).

In the specimens of this paper, some of the extruded papillae, also the papillae in histological sections present two lobes, in some cases each with own duct. Especially interesting is the different aspect of the ejaculatory duct in specimens Fl1 and Fl2: very wide and filled with sperm in specimen Fl1; with a possible narrow and convoluted course in specimen Fl2. At my current state of knowledge, it was not established when and how / by what mechanisms the ejaculatory duct is carved inside the penis tissue.

#### 3) The atrial cavities

The interpretation of the small atrial cavity (common atrium or the vaginal area of the bursal canal) has a very much phylogenetic weight.

a) in frontal sections, because of the section plan, the existence of a vaginal part of the bursal canal or that of a common atrium is difficult to be established. The small ventral cavity the atrial tube enters can be considered either as a common atrium, as well as the vaginal area of the bursal canal. It is important to be noticed that the oviducts open separately in this small ventral cavity, very close to the atrial tube.

b) in sagittal sections, the slides Fl1-27-2 and Fl1-27-4 (Figs. 5, 6) show the entry of the atrial tube into a distinct cavity positioned ventrally. The slide Fl1-26-1 (Fig. 4) shows what appears to be the entry of the bursal canal into a ventral cavity communicating with the exterior through the gonopore. These sections demonstrate the existence of two distinct cavities: a common atrium, different that of the terminal part of the bursal canal (the vaginal area of the bursal canal). The gonopore should be considered the orifice by which an atrium (and not the bursal canal) opens to the exterior.

Given the oviducts (at least one oviduct) open very close to the atrial tube – Figs. 15, 16, it implies that the oviducts open into a common atrium. If these oviduct openings are real / correctly interpreted, this would have consequences on the phylogeny of the higher taxa, being known that in Dugesiidae the oviducts open into the bursal canal while the openings of the oviducts into the common atrium is characteristic for Planariidae and Dendrocoelidae (DeVries & Sluys 1991, Sluys 2001).

The openings of the oviducts, the atrial cavities and their communications in general deserve more attention, especially the literature present various kind of ducts – “the Drüsengand, the canalis anonymus, the Beauchamp’s canal”, even if in the situation of Terricola (Sluys 2001).

#### 4) The atrial fold

A well-developed atrial fold is typical for *S. mediterranea* (Benazzi et al. 1975, Leria et al. 2018), yet, not mentioned for the Tunisian populations (Harrath et al. 2004). An atrial fold has rarely been observed in *S. polychroa* and *S. lugubris* (Benazzi et al. 1970). It is absent in *S. nova*, also in the specimens of this paper.

#### 5) A nipple on the tip of the penis papilla

This character is typical for *S. lugubris*, as a permanent character (Reynoldson & Bellamy 1970, Ball & Reynoldson 1981, Reynoldson & Young 2000, Leria et al. 2018). An eversible nipple-like tip can be seen in *S. polychroa* (Reynoldson & Bellamy 1970) in living specimens. It is absent in *S. mediterranea*. Some specimens of *S. nova* show a nipple on the tip of the penis papilla (Leria et al. 2018). A sort of a large nipple is visible in specimen Fl1 of this paper – Fig. 7.

The Schmidtea population presented in this paper:

1. shows similarities with other Schmidtea species:
  - with *S. polychroa*, the presence of sensory fossae, as presented by DeVries & Sluys (1991)
  - with *S. mediterranea*, the reproduction by fission (Riutort et al 2012)
  - with *S. nova*, a diaphragm of the ejaculatory duct to the secondary seminal vesicle (Leria et al. 2018)
2. shows its own characters:
  - the disposition of the genital atria
  - the wide ejaculatory duct inside the penis papilla
  - at least some specimens with a bilobed penis papilla 3) does not fit the diagnosis / characteristics of any other Schmidtea species, thus, it is not any of the other Schmidtea species because – see Table 1

**Table 1.** Summary comparison of Schmidtea species.

| Schmidtea species | Character | Schmidtea population of this paper |
| --- | --- | --- |
| *S. lugubris* | conic penis papilla | bilobed papilla |
| *S. polychroa* | conic penis papilla |  |
| *S. mediterranea* | conic penis papilla |  |
| *S. nova* | penis papilla knee-bend shape |  |
| *S. mediterranea* | atrial fold | atrial fold absent |
| *S. nova* | egg-shaped formation of the bursal canal | egg-shaped formation of the bursal canal absent |
| *S. nova* | interconnected testis follicles, parovaria (Leria et al. 2008) | not observed |
| *S. nova* | sclerotized/chitinized wall of the distal ejaculatory duct (Leria et al. 2004) | not observed |

## Abbreviations used

at: atrial tube
bc: bursal canal
ca/vg: the common atrium/the vaginal area of the bursal canal
cb: copulatory bursa
ej: ejaculatory duct
go: gonopore
ma: male atrium
nip: nipple
ovd: oviduct
pb: penis bulb
pp: penis papilla
psv: primary seminal vesicle
sh: shell glands
spv: spermiducal vesicles
ssv: secondary seminal vesicle
sw: swelling
vd: vas deferens

